# Sex-, development-, and nutrition-dependent expression of a major age-at-maturity gene in Atlantic salmon

**DOI:** 10.1101/2024.09.15.613130

**Authors:** Eirik Ryvoll Åsheim, Paul Vincent Debes, Andrew House, Petra Liljeström, Annukka Ruokolainen, Morgane Frapin, Iikki Donner, Ehsan Pashay Ahi, Jaakko Erkinaro, Jukka-Pekka Verta, Craig R Primmer

**Author notes:** Authors contributed equally (Jukka-Pekka Verta, Craig R Primmer).

## Abstract

Sexual maturation is a key process in the life history of an organism and is influenced by both genetic and environmental factors. In Atlantic salmon (Salmo salar), alternative alleles at the gene vgll3 have been found to associate with a large proportion of the variation in maturation age, yet the molecular and physiological mechanisms of this association are still largely unknown. Here, we investigated the mechanisms by which genetic variation in this major age-at-maturity gene translated into phenotypic variation. We established a time series of gonadal vgll3 mRNA expression in male and female Atlantic salmon in a common garden setting, starting at the prepubertal stage and spanning multiple spawning seasons. We found that vgll3 mRNA expression is reduced in early-maturing males and females. However, for the second spawning season, vgll3 expression increased with gonadal development in males, but eventually decreased in females. Expression of vgll3 had context-dependent correlations with expression of amh and lhr. We also found signs of reduced testicular vgll3 expression in a low-fat feed treatment. By significantly widening the scope of our knowledge on temporal vgll3 expression dynamics, our results exemplify that major-effect genes may use different pathways in sexes but achieve similar outcomes.

## INTRODUCTION

Sexual maturation is a process that spans a considerable period of an organism’s growth and development. This process is also tightly linked with the allocation of resources to reproduction, thus having a central role in determining organismal life history and ecology (Hutchings, 2021). Variations in the maturation process manifest themselves in different forms, one of which is the age at maturity – the amount of time an organism spends from its conception until it is reproductively mature. Atlantic salmon display remarkable variation in age at maturity, ranging from one to ten years (Hutchings & Jones, 1998). In European populations, a large fraction of this variation in age at maturity has been linked to variation at a single genetic locus – *vgll3 –* for both males and females (Barson et al., 2015). Although such large-effect loci have been discovered for a range of life-history traits in various organisms (Johnston et al., 2013; Küpper et al., 2016; Lamichhaney et al., 2016; Pearse et al., 2019; Troth et al., 2018), knowledge on the functional connections (molecular and physiological) between their genotypes and respective phenotypes is largely lacking. Consequently, much remains to be learned about the broader implications of these genes, including other phenotypic effects, environmental interactions, and maintenance of genetic variation. The link between *vgll3* and age at maturity is a notable exception to this pattern, as several facets of the processes connecting *vgll3* genotype and age at maturity phenotype have been uncovered since its initial discovery.

In Atlantic salmon, two alleles of *vgll3* associate with either early (*vgll3*\*E) or late (*vgll3*\*L) maturation. This association was initially observed in wild European Atlantic salmon males and females (Barson et al., 2015), and has subsequently been confirmed in multiple laboratory common-garden studies (Åsheim et al., 2023; Debes et al., 2021; Sinclair-Waters et al., 2022; Verta et al., 2020). Furthermore, *vgll3* has been shown to have pleiotropic phenotypic effects, with alternative genotypes being associated with body condition (Debes et al., 2021; House et al., 2023), aerobic scope (Prokkola et al., 2021), as well as aggressive behaviour (Bangura et al., 2022). At the molecular level, *vgll3* genotype has been associated with differing expression of genes relating to the HPG axis in the pituitary and in the testis (Ahi et al., 2022, 2023), and *vgll3* alleles have been associated with alternative splicing (Verta et al., 2020). Correlated RNA expression of other genes has indicated that *vgll3* might be active in gene networks regulating cell differentiation and proliferation, such as the Hippo pathway (Kjærner-Semb et al., 2018; Kurko et al., 2020). The timing and location of *vgll3* expression itself has indicated an important and direct role in testis development (Kjærner-Semb et al., 2018; Verta et al., 2020). Interestingly, *vgll3* has been found to have similar associations in a range of other vertebrates. In humans, genetic variation at- or close to the *vgll3* locus has been associated with age at maturity (Cousminer et al., 2013; Elks et al., 2010; Perry et al., 2014), adiposity-related traits (Nakayama et al., 2017), cancer (Hélias-Rodzewicz et al., 2010), and autoimmune disease (Liang et al., 2017). In mice, *Vgll3* has been identified as an inhibitor of adipocyte differentiation (Halperin et al., 2013), and is involved in muscle development (Figeac et al., 2019; Mielcarek et al., 2009) and testis development (McDowell et al., 2012). Overall, this strongly suggests a conserved function of *vgll3* across vertebrates, but also shows how *vgll3* seems to have a role that is active in multiple cellular processes, making it hard to pinpoint a single “starting point” for the influence of *vgll3* on age at maturity.

Gonadal tissues and in particular Sertoli and granulosa cells have been identified as key sites of *vgll3* expression in Atlantic salmon testis and ovaries (Kjærner-Semb et al. (2018)). Expression of *vgll3* was also found to be higher in immature-than in mature testes (Kjærner-Semb et al. (2018), Verta et al. (2020)). This has indicated that *vgll3* may function as an inhibitor of testis development. Specifically, based on the down-regulation of *vgll3* and Hippo pathway members in pubertal testis, Kjærner-Semb et al. (2018) hypothesized that *vgll3* controls the early Sertoli cell proliferation which is necessary for the further development of spermatogonial cells (França et al., 2015; Schulz et al., 2010). Significant knowledge gaps remain regarding how *vgll3* is expressed in gonads during the maturation process. For example, earlier studies only cover early-spawning males and no mature females. Thus, *vgll3* expression in relation to gonadal maturation for later-maturing males and females is not known. Further, the data available for *vgll3* RNA expression in wild-derived Atlantic salmon covers only early parr maturation (1^st^-year maturation, Kurko et al., 2020; Verta et al., 2020). Some data are also available for the post smolt stage (Kjærner-Semb et al., 2018), though only from aquaculture strain Atlantic salmon. Finally, some studies have suggested that the influence of *vgll3* on age at maturity might be mediated via effects on adiposity and resource acquisition (Debes et al., 2021; Halperin et al., 2013; House et al., 2023), but there have been no studies so far looking at how nutrient availability (or other environmental factors) influence *vgll3* RNA expression.

Here, we sought to expand the knowledge of the role of *vgll3* during the maturation process by quantifying its gonadal mRNA expression over multiple years in a single experimental cohort. The time series presented here covers gonadal *vgll3* expression levels in wild-derived Atlantic salmon, starting at the immature stage (second spring, two years old) and spanning multiple subsequent spawning seasons for both female and male individuals (up to four years old). This study also included a feed nutrition treatment designed to test if the influence of *vgll3* varies with environmental variation in nutrient availability. We found patterns of expression supporting previous findings of gonadal *vgll3* expression being associated with delayed maturation in males and found the same pattern for early-maturing females. Unexpectedly, we found striking differences in patterns of expression over gonadal development for third- and fourth-year spawners. These results suggest a different mode of action in testes and ovaries and provides a basis for development of new hypotheses and research on the mechanistic pathways of *vgll3* associated with maturation.

## METHODS

### Experimental design

This study used Atlantic salmon (*Salmo salar*, L. 1758) from the same rearing experiment as described in Åsheim et al. (2023). In brief, parents with known homozygous *vgll3* genotypes were crossed in late October 2017 to create a series of 2 × 2 factorials (one of each sex of each homozygous *vgll3* genotype), resulting in one family with all offspring having the *vgll3*EE* genotype, one family all *vgll3*LL* and two families all *vgll3*EL*. In total, 17 factorials were created using 67 parental individuals, resulting in 68 unique crosses (Figure 1). At first feeding, the fish were divided evenly (for families) among 12 circular flow-through tanks (1.00 m high, 2.77 m wide, up to 5.4 m^3^) at Lammi Biological Research Station (University of Helsinki, Lammi, Finland). Water was pumped directly from the local lake Pääjärvi, and thus the water temperatures in the tanks closely followed the daily and seasonal lake water variation. The 12 tanks were divided among two temperature treatments, using a heat exchange system to divide the incoming water into a cold and a warm temperature treatment (2°C difference). The current study only used fish from the warm treatment, thus using fish from six of the 12 tanks, which experienced temperatures following those of the lake plus approximately 1°C. Note that in March 2021, after completion of the experiment including both the warm and cold temperature treatments, the cold-temperature tanks were cleared out and individuals from each of the six warm-treatment tanks were divided among 12 tanks for the final year of the experiment (Figure 1 B).

**Figure 1.**
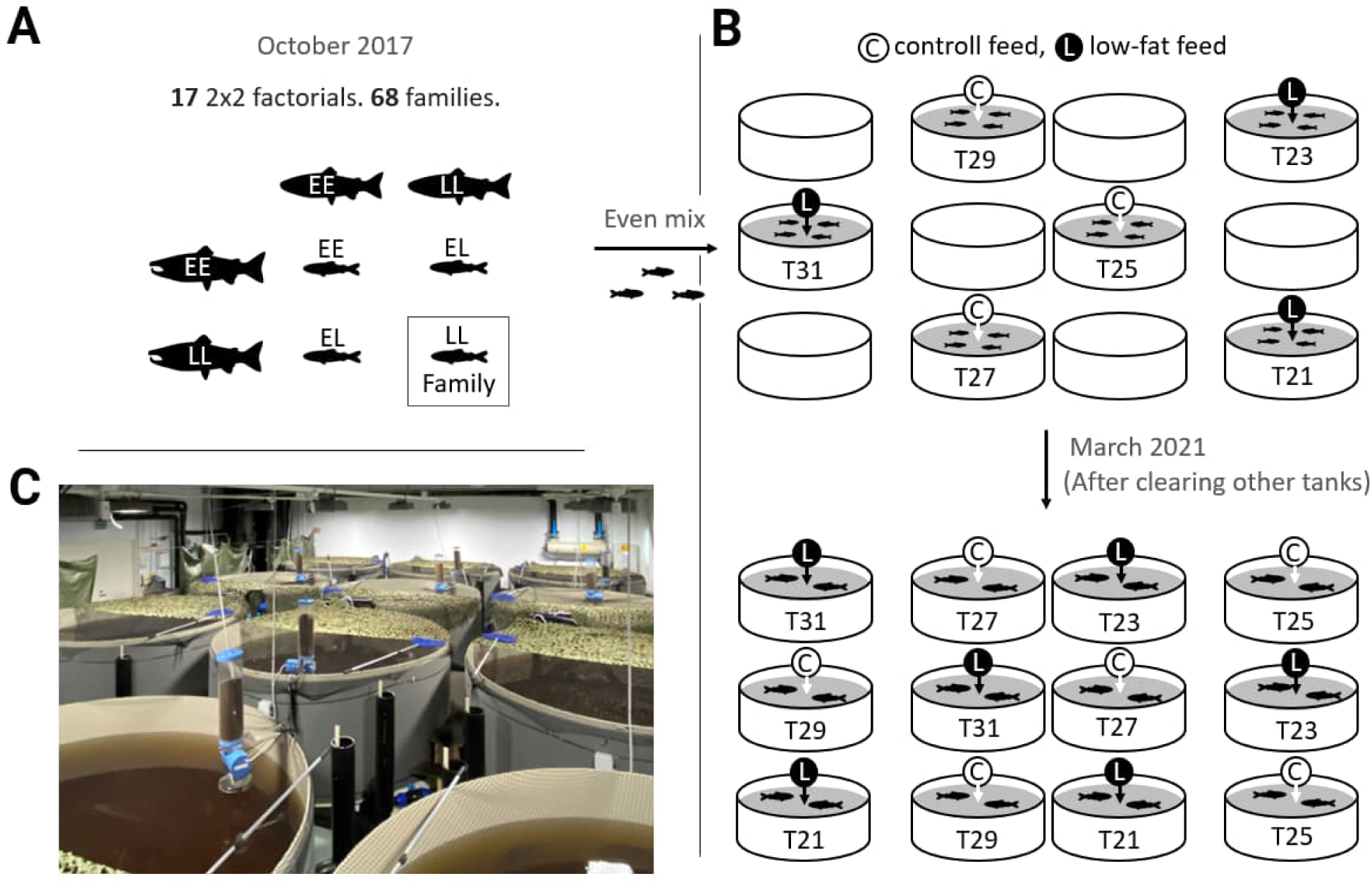
Experimental design. Shows the crossing design (A) the allocation of fish used in this study to different tanks and treatments (B), and a photograph (C) of the experimental system (taken in March 2020).

To study potential gene-environment interactions between *vgll3* genotype and nutrition, tanks were subjected to one of two feeding treatments, “control” and “low-fat”. At the beginning of the study (3/2018), all fish were fed ad-libitum using body-size-matched pellets of commercial fish feed (”Hercules”, Raisioaqua, Raisio, Finland). In July 2019 (summer of the second year post-fertilization), the feed for half of the tanks was changed to a custom-made fat-reduced feed of the same brand and pellet type. Thus, from July 2019, fish were either fed the control feed (17-26% fat, 18.10-20.40 kj g^−1^ depending on pellet size), or the low-fat feed (12-13% fat, 17.25 kj g^−1^).

To keep track of fish identities, all fish were marked with a passive integrated transponder tag (PIT tag) at the first measurement in April 2019. Tags were inserted into the abdominal cavity, slightly behind (posterior to) their right-side pectoral fin. Additionally, a small fin clip was taken from their caudal fin for genotyping, sex determination, parental assignment and *vgll3* genotype confirmation (as in Debes et al., 2021).

Fish were reared until March 2022, after which almost all male and female fish had matured. At this point, all fish were euthanized by anaesthetic overdose (MS-222, Tricaine methanesulfonate, 0.250 g L^−1^, sodium bicarbonate buffered).

The fish used in this study were offspring of first-generation hatchery fish. The parental broodstock is used for supplementary stocking of Finnish rivers and has its name “Neva” from the river where it was mainly sourced in the 1980s. Since that time, it is renewed every few years using fish that have completed a seaward migration after their release. Thus, the grandparents of our source experimental population had completed a full marine migration, while all the parents had experienced the same environment in the hatchery (thus minimizing potential maternal/epigenetic variation, such as the environmental variation experienced by migrating individuals).

### Tissue sampling

Fish were measured and sampled four times per year, starting in April 2019 and continuing until the end of the study in March 2022, resulting in 13 time points (see Table 1 for an overview). At these time points, phenotypic traits were recorded for each individual (details below). For each tank, eight individuals were euthanized for tissue sampling one day after measurements, except for time point 3, where they were sampled seven to 12 days after, and time point 4, where they were sampled on the same day. Measuring and sampling at each time point was conducted over a period of 14-16 days.

**Table 1.** Sample overview. Overview of tissue-sampled individuals that were included in this study. Sample sizes are shown per sex and *vgll3* genotype at each time point. Note that this is a subset of the total amount of sampled fish, as both heterozygous *vgll3* genotypes were sampled (both EL and LE) from each tank (n=6) and sex (n=2), but only half were included in the analysis (randomly selected). No fish were sampled for tissues at time point 1 (only measurements).

| Time point | Year | Month | n | Male | EE | EL | LL | Female | EE | EL | LL |
| --- | --- | --- | --- | --- | --- | --- | --- | --- | --- | --- | --- |
| 2 | 2019 | July | 2 | 11 | 3 | 4 | 4 | 14 | 6 | 5 | 3 |
| 3 | 2019 | October | 3 | 18 | 5 | 6 | 7 | 17 | 5 | 6 | 6 |
| 4 | 2019 | November | 3 | 18 | 6 | 6 | 6 | 18 | 6 | 6 | 6 |
| 5 | 2020 | February | 3 | 17 | 5 | 6 | 6 | 19 | 6 | 6 | 7 |
| 6 | 2020 | May | 3 | 16 | 5 | 6 | 5 | 17 | 5 | 6 | 6 |
| 7 | 2020 | August | 3 | 19 | 7 | 6 | 6 | 19 | 7 | 6 | 6 |
| 8 | 2020 | November | 3 | 15 | 5 | 5 | 5 | 17 | 6 | 6 | 5 |
| 9 | 2021 | February | 3 | 16 | 5 | 6 | 5 | 17 | 4 | 6 | 7 |
| 10 | 2021 | May | 3 | 20 | 6 | 8 | 6 | 19 | 5 | 9 | 5 |
| 11 | 2021 | August | 3 | 17 | 6 | 6 | 5 | 19 | 6 | 7 | 6 |
| 12 | 2021 | November | 3 | 18 | 6 | 6 | 6 | 17 | 6 | 6 | 5 |
| 13 | 2022 | February | 3 | 19 | 6 | 7 | 6 | 17 | 6 | 6 | 5 |

Phenotypes recorded were body mass, body length, maturation status, and whether fish showed external signs of smoltification (migration phenotype). Male maturation status was checked by stroking the abdomen towards the vent, recording any fish releasing milt as mature. Females were recorded as mature if releasing eggs. Migration phenotype, being either migrant (smolt) or resident (parr), was scored based on silvering of color and loss of parr marks. Individuals were recorded as having smolted from the time point following the last recording of resident (parr) characteristics. For measurements, fish were netted out of their tank and placed in an aerated anaesthetic bath (MS-222, 0.125 g L^−1^, sodium bicarbonate buffered) within 1°C of the tank temperature. After fish stopped responding to touching or holding, their weights were measured down to the nearest 0.01 g (April and July 2019) and subsequently 0.1 g using a digital scale (Scout STX222 or STX6201, Ohaus, Parsippany, USA). Fork length (snout to fork of tail) was measured using a digital fish-measurement board (DCS5, Big Fin Scientific, Austin, TX, USA). Finally, migration and maturation phenotypes were recorded before the fish were returned to their tank.

A subset of fish was selected for tissue sampling at each measurement time point. From each tank, one individual of each sex-*vgll3* genotype combination was selected for sampling, avoiding selecting multiple fish from the same family within tanks, totalling 2 × 4 × 6 = 48 sampled individuals per measurement period. At the first sampling time point (time point 2, Table 1), the sex and genotypes of the fish were unknown, so 10 fish were randomly sampled from each tank. For all time points following time point 4, fish selected for sampling were placed directly into a separate holding tank after measurements and were taken from this holding tank for sampling on the following day. On the preceding time points (2, 3 and 4), fish to be sampled were netted from their original holding tank.

At sampling, fish were first euthanized using an MS-222 overdose (0.250 g L^−1^, sodium bicarbonate buffered), and were then immediately dissected for collection of tissue samples. Sampled gonads were immediately placed in 1.5 ml Eppendorf tubes and flash-frozen in liquid nitrogen before being stored at −80°C. Gonads were sampled whole, and in separate tubes for as long as their size allowed this. For gonads that were too large to fit the 1.5 ml Eppendorf tubes, only the tip of the gonad (anterior end) was sampled. Mature gonads were sampled without stripping eggs or sperm, but sperm was removed from samples before RNA extraction.

### RNA extraction and quantification

Before RNA extraction, testis samples were stripped for sperm by thawing them in room-temperature phosphate-buffered saline (PBS) for ∼1 minute and then squeezing and rolling the tissues on a piece of paper tissue (coffee filter paper) until only solid tissue remained. This procedure was done on all testis samples regardless of maturation status and lasted 2-3 minutes from start of thawing.

To extract total RNA, we used the NucleoSpin 96 RNA kit (MACHEREY-NAGEL, Dueren, Germany). Samples were homogenized in RA1 buffer with added DTT (final concentration 10 mM) using a Bead Ruptor Elite (Omni Inc, Kennesaw, GA), 2 ml or 7 ml tubes, and 2.4 mm steel beads. Residual DNA was removed with TURBO DNase (Invitrogen, Waltham, MA). To avoid any batch effects confounding other variables, samples were randomly divided among the five plates used for RNA extraction and subsequent RT-ddPCR (reverse transcription digital droplet PCR).

We measured RNA concentrations using ThermoFisher Quant-it reagents and diluted the samples to a 10 ng/µl working concentration. Dilution concentrations were measured with ThermoFisher Qubit High Sensitivity reagents and were re-adjusted if needed. Final concentrations were verified with Qubit HS. To quantify the absolute level of expression of *vgll3* (chromosome 25), *amh (*anti-müllerian hormone), and *lhr* (luteinizing hormone receptor), we used one-step RT-ddPCR (BioRad) and TaqMan probes (Supplementary table 1). For allele-specific quantification of *vgll3*, we used TaqMan probes specific to the *Early* and *Late vgll3* alleles as in Verta et al. (2020). We used ∼40 ng of total RNA as template for *vgll3*, and ∼20 ng for *amh* and *lhr*.

To extract counts of detected mRNA molecules, RT-ddPCR results were analysed in *BioRad Quantasoft* v.1.7.4.0917. For most samples, we used an automatic selection of signal thresholds, and used manually set thresholds for samples where the software was not able to select them automatically. All 1-D amplitude plots were visually checked, and threshold settings verified. We excluded samples with less than 5000 accepted droplets (unreliable estimates) as well as samples with zero negative droplets (extreme values). Finally, we normalized the counts of detected *mRNA* molecules to the amount of total template RNA so that mRNA expression for *vgll3, amh, and lhr* was expressed as number of copies per ng of total RNA.

mRNA counts for ovaries were corrected for 5S rRNA amounts since the amount of this ribosomal RNA changes during development and can have a significant influence on the total RNA count (Shen et al., 2017). This was corrected by determining the proportion of 5S rRNA from electropherograms acquired using a 2100 Agilent Bioanalyzer instrument, and then subtracting this amount from the total RNA before calculating the number of copies (*vgll3*, *lhr*, *amh*) per ng RNA. Thus, expression for ovaries is expressed as total RNA excluding 5S rRNA.

### Overview of animals and samples

The experiment was conducted under an animal experiment permit granted by the Finnish Project Authorization Board (Permit ESAVI/2778/2018)

Over the entire study period, 562 fish were tissue-sampled. The 417 individuals analysed for this study are a subsample of this due to only half of the *vgll3* heterozygotes being included. Heterozygotes were randomly selected for each combination of tank, time-point, and sex (Table 1). For some statistical tests, the sample size was smaller than the original number of individuals included due to missing data. Overall, the number of individuals with missing mRNA expression data was 17 for *vgll3*, 27 for *lhr*, and 49 for *amh*.

In 2019, the starting number of Neva individuals in the warm-temperature treatment (the subset of the source experimental population used here) was 1162 individuals, and 411 remained at the conclusion of the experiment in 2022. Of the 751 fish which died throughout the experiment, 255 (22%) were unplanned mortalities which were not part of any sampling or other procedure.

Figure 2 shows the number of individuals in different stages of maturation for each time point. Most sampled individuals had undergone smoltification. Exceptions were for two, two and three females in time points 2, 3 and 4, respectively, and two, three and one males in time points 3, 4 and 10, respectively having parr characteristics.

**Figure 2.**
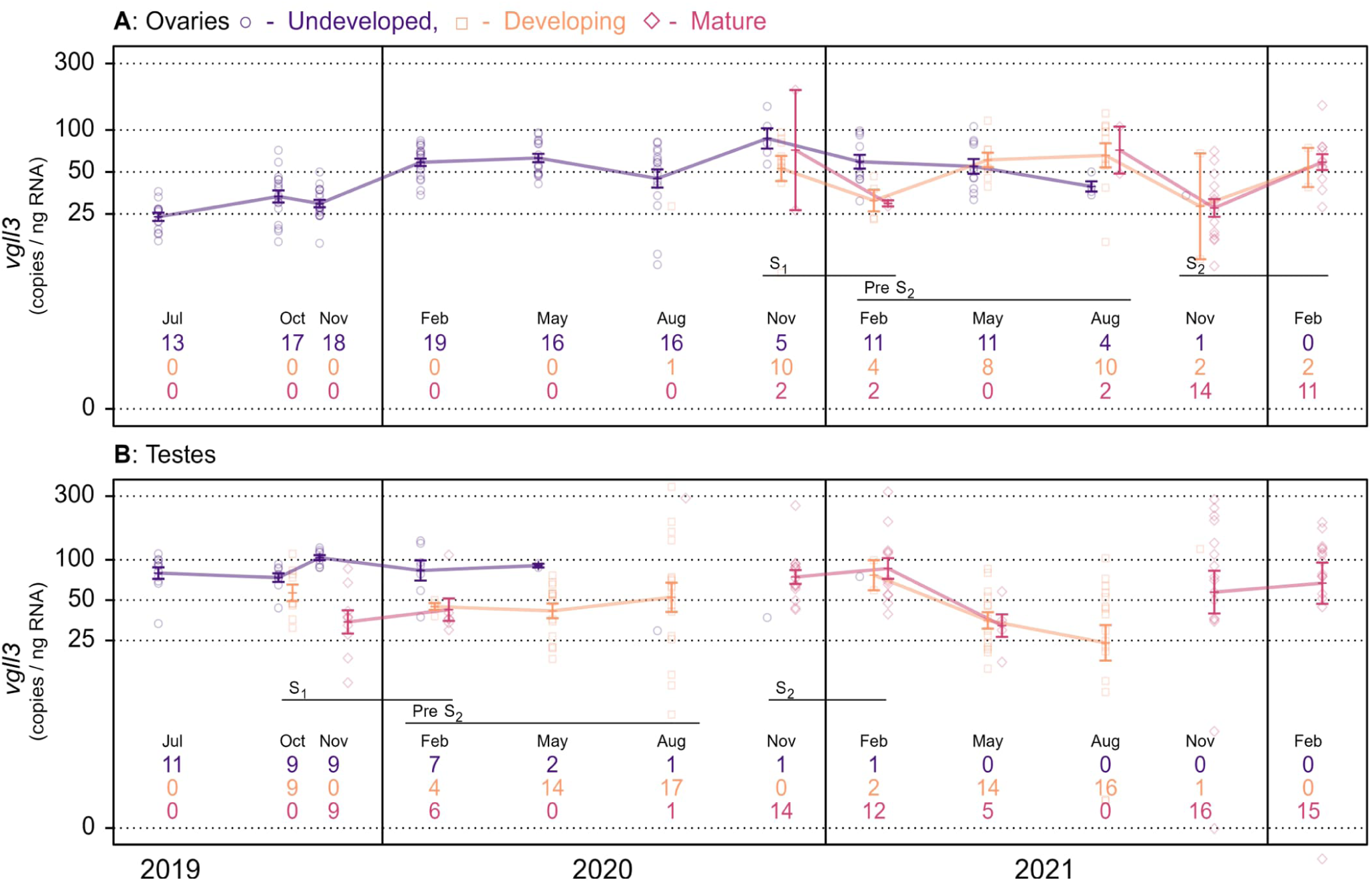
Gonadal maturation and *vgll3* expression over time. Gonadal expression of *vgll3* mRNA over the entire study period, measured as copies ng_-1_ of RNA in ovaries (A) and testes (B). Colours and shapes correspond to maturation status of the gonads: purple circles for undeveloped gonads (no sign of enlargement), orange squares for developing gonads (some sign of enlargement), and red diamonds for completely mature gonads (releasing sperm or eggs). Lines connect means between time points with at least two samples, and error bars indicate SE. Numbers below points indicate sample sizes for undeveloped, developing, and mature gonads. The y-axes are log_e_ scaled.

### Statistical methods

Our statistical analysis approach aimed to test for *vgll3* mRNA expression differences between fish at different developmental stages and to explore seasonal or developmental patterns in expression of *vgll3* mRNA (and associations with *vgll3* genotype). Additionally, having access to mRNA expression levels of *amh* and *lhr*, we explored potential correlations between *vgll3* and *lhr* /*amh* expression. Finally, we tested if the association between *vgll3* genotype and age at maturity was detectable in our sample of individuals. For all comparisons via models (except correlations), we included feeding treatment as an explanatory factor to assess if this affected any of the expression patterns.

Males and females started spawning in different years. To simplify comparisons between sexes, we analysed the data relative to the first spawning season. Thus, we refer to the first and second spawning seasons, which were in 2019 and 2020 for males, and 2020 and 2021 for females (Figure 2, lines S1 and S2).

### Description of analysis

Gonads were classified into one of three stages of development: 1) “undeveloped” (immature) testes were identified as those resembling a thin transparent string running dorsally along the entire length of the body cavity. In females, we identified undeveloped ovaries as those having a length less than 1/4^th^ of the body cavity. 2). “Developing” testes were classified as those being thicker than the undeveloped ones and having a white or grey hue. Developing ovaries were identified as those being longer than 1/4^th^ of the body cavity. 3) “Mature” testes and ovaries were identified as those releasing eggs or sperm directly from the fish. Figure 2 shows the number of fish in each category for each sampling time point.

For all models described below, the IDs of each individual’s dam, sire, and family were included as random effects to account for parental- and family effects. Tank ID was also included as a random effect to account for potential tank effects. Males and females were analysed in separate models.

Model 1: Here we tested for differences in *vgll3* expression between maturation stage and feed treatment in the first spawning season. We simplified the grouping of the gonads into two categories, being either undeveloped or developing/mature. We then specified models testing if the log of standardized *vgll3* mRNA expression (counts ng^−1^ RNA) changed with maturation status, feeding treatment, or month of sampling (November or February). Interactions between feed and both maturation status and month of sampling were also included.

Model 2: Here we tested the observed increase in *vgll3* expression in the time point leading up to the second spawning season in developing gonads (Figure 2, lines “Pre S2”). To do this, we specified models testing if the log of standardized *vgll3* expression (counts ng^−1^ RNA) changed with month of sampling (2, 5, 8), feeding treatment, and whether the sampled individual had been recorded as mature on a previous occasion. Interactions between month of sampling and both feed and previous maturation status and were also included.

Model 3: To test for differences in maturation probability in the first spawning season between *vgll3* genotypes and feed treatments, we used generalized linear mixed effect models with Bernoulli error distribution and logit link function. Maturation was categorized in the same two categories as in model 1, and the model tested if maturation probability differed between *vgll3* genotypes and feed treatments. The body weight of each individual in the proceeding spring, the time point where growth is expected to have its most significant effect on future maturation (Rowe et al 1991), was included in the model to account for body-size differences. Interactions between feed and both maturation status and month of sampling were also included.

Correlations: Expression correlations between *vgll3* and *amh* or *lhr* were estimated using Pearson correlations on log-adjusted standardized expression levels (counts ng^−1^ RNA). Correlations were estimated separately on data grouped by sex, maturation status, and season of the year (spring, summer, autumn, winter). In the final month before the second spawning season, which was the time point where we observed the largest variance in gonadosomatic index (GSI) in developing testes and ovaries, we checked whether this variation was associated with expression of *vgll3* using Pearson correlations between log gene expression and GSI.

To verify that the feeding treatment had an impact on fish growth, we performed *t*-tests on body condition between fish in the control- and low-fat feed treatment for each measurement time point. Body condition was calculated as body mass (g) divided by the cube of the body length (cm) multiplied by 10 000 (Fulton’s K)(Ricker, 1975). This analysis confirmed that body conditions between the two treatments were significantly different at every time point after May 2020 (the low-fat feed treatment started in July 2019) (Supplementary figure 3).

For all gene expression level analyses, we log-transformed expression level of *vgll3*, *amh*, and *lhr* (copies ng^−1^ RNA) to account for heteroscedasticity (higher variance at higher expression levels). GSI was calculated as percent gonad mass of total body mass.

### Technical model descriptions

The following formulae describe the models introduced above, showing the overall form of the models, specifically which variables were included as explanatory and response variables, and which interactions were included. Each model was fitted separately for each sex (testes, ovaries). Not shown: parameters *b*_*i*_ for each term (*i*) of the models (except *R* and *e*).

Model 1: *vgll3* expression differences between maturation stages in the first spawning season.

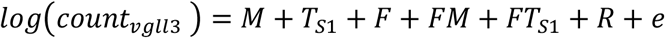

Model 2: Gradual change in *vgll3* expression leading up to the second spawning season.

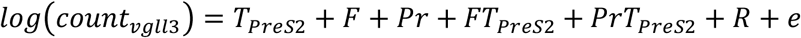

Model 3: Effects on maturation probability

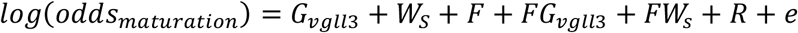

| Variable | Explanation |
| --- | --- |
| $count_{vgll3}$ | *vgll3* expression (mean-centred, SD-scaled log of expression, copies ng<sup>-1</sup> RNA). |
| $odds_M$ | Odds of gonads having started maturing or not. |
| $M$ | Gonad development having started or not (0=no, 1=yes). |
| $T_{S1}, T_{PreS2}$ | Month of sampling, during the first spawning season (S1: 0=Dec, 1=Feb), or leading up to the second spawning season (PreS2: 0=Feb, 3=May, 6=Aug). |
| $F$ | Feed treatment (0=low fat, 1=control). |
| $Pr$ | Whether the sampled individual had been recorded as mature earlier or not (0=not previously mature, 1= previously mature). |
| $G_{vgll3}$ | *vgll3* genotype (LL=-1, EL=0, EE =1). |
| $W_s$ | Body weight in the preceding spring (mean-centred, SD-scaled log of weight). |
| $R + e$ | The random effects on the intercept (dam ID, sire ID, their interaction, and Tank ID) and residual error. |

Bayesian model parameters (b_i_, R and *e*) were estimated using *stan*, which was interfaced via the *R* package *brms*. All models were run for 5 000 iterations in four chains, discarding the first 500 iterations as warmup, resulting in 18 000 posterior samples of each parameter. All model fits were verified using a visual posterior predictive check (function *pp_check* in R package *brms*) and verified for influential points by inspecting pareto k diagnostic values (function *loo* in R package *loo*) (Vehtari et al., 2017). Three models had high numbers of influential data points (points for which k > 0.7), which were male model 2 (3 of 42 points with k > 0.7), female model 2 (6 of 26 points with k > 0.7), and male model 1 (8 of 35 points with k > 0.7). For these models, we tested alternative versions without random effects for parental- and family ID, which reduced the number of influential points down to 0, 2, and 1, respectively, indicating that most of these influential points were affecting the estimates of parental- and family effects. We compared effect-size estimates of the models including- and excluding parental- and family effects, found that the differences were small, and chose to keep the models including parental- and family effects.

We chose relatively non-informative priors for all parameters. We used normally distributed priors with a mean of 0 and SD of 1 for the intercept and for all effect-size parameters. For error terms, we used normally distributed priors with a mean of 0.2 and SD of 0.5.

A summary of the models’ fixed-effect parameter estimates (*b_i_*) is presented in Supplementary Figure 1. Where interactions were non-significant (95% credible intervals including zero), we simplified parameter estimates by calculating unconditional (marginal) estimates of the main effects (Supplementary figure 1).

All statistical analysis was done in the *Rstudio* v2022.7.2.576 (RStudio Team, 2020) software environment running *R* v4.2.2 (R Core Team, 2021) and *Rstan* v2.21.8 (Stan Development Team, 2022) R packages *brms* v2.18.0 (Bürkner, 2017) was used for creating *Rstan* models, *loo* v2.5.1 (Vehtari et al., 2022) for inspecting pareto k diagnostic values, *bayesplot* v1.10.0 (Gabry & Mahr, 2022) for posterior predictive checks, *ggplot2* v3.4.0 (Wickham, 2016) for producing plots, and *tidyverse* v1.3.2 (Wickham et al., 2019) for data workflow.

## RESULTS

### Expression of *vgll3* across 3 years of development

We measured the gonadal expression of *vgll3* mRNA (copies ng^−1^ RNA) in different seasons over multiple years, in both male and female Atlantic salmon. We found that *vgll3* expression was elevated in both undeveloped testes and ovaries in the first spawning season, but also that expression patterns were more complex for the following spawning seasons.

In the first spawning season, in both testes and ovaries (Figure 2, lines S1), gonadal *vgll3* expression was higher in undeveloped gonads than in those showing signs of maturation (i.e., gonads enlarged or fully mature). For ovaries, *vgll3* expression was 1.59 times higher [1.05, 2.34] (brackets indicate 95% credible intervals) in undeveloped than in developing or mature ovaries, with a predicted mean of 44.12 copies ng^−1^ RNA in developing or mature ovaries, and 69.88 copies ng^−1^ RNA in undeveloped ovaries (posterior predictions). For testes, *vgll3* expression was 2.05 times higher [1.44, 2.89] in undeveloped testes (same comparison), with a predicted mean of 40.05 copies ng^−1^ RNA for developing or mature testes, and 81.98 copies ng^−1^ RNA in undeveloped testes.

In the second spawning season (Figure 2, lines S2), we were unable to directly compare maturing and undeveloped individuals as most individuals were maturing. However, looking at the period leading up to the second spawning season (Figure 2, lines Pre S2), there was a trend of increasing *vgll3* expression up towards spawning. In testis, *vgll3* expression increased 1.20-fold [1.21, 1.40] for each month going from February (39.22 copies ng^−1^ RNA) to August 2020 (86.46 copies ng^−1^ RNA). For females, ovary *vgll3* expression increased 1.12-fold [1.02, 1.24] at each time point from February (33.1 copies ng^−1^ RNA) to May to August 2021 (69.8 copies ng^−1^ RNA). After this period, as maturation occurred, testis *vgll3* expression remained high, while ovary *vgll3* expression first dropped in November, but then increased again in the following February (Figure 2, lines S2). Thus, *vgll3* expression increased as testes and ovaries were developing, which contrasts with the first season where the undeveloped gonads had the highest *vgll3* expression. Notably, in testes, this pattern was dependent on feed treatment and was only present in the control feed treatment (detailed below). Whether fish had matured in the first spawning season had no detectable effect on *vgll3* expression in the period leading up to the second spawning season. In this analysis, 17 of 42 male individuals- and 9 of 26 female individuals had matured previously.

GSI and *vgll3* correlated significantly at the last time-point before the second spawning season, which was when GSI had been increasing the most (August, Figure 3 A and B). However, this correlation was sex dependent. In testes, a higher GSI corresponded to a higher expression of *vgll3* (Pearson’s R = 0.79, p < 0.001), while expression of *vgll3* decreased with GSI in ovaries (R =-0.92, p = 0.001).

**Figure 3.**
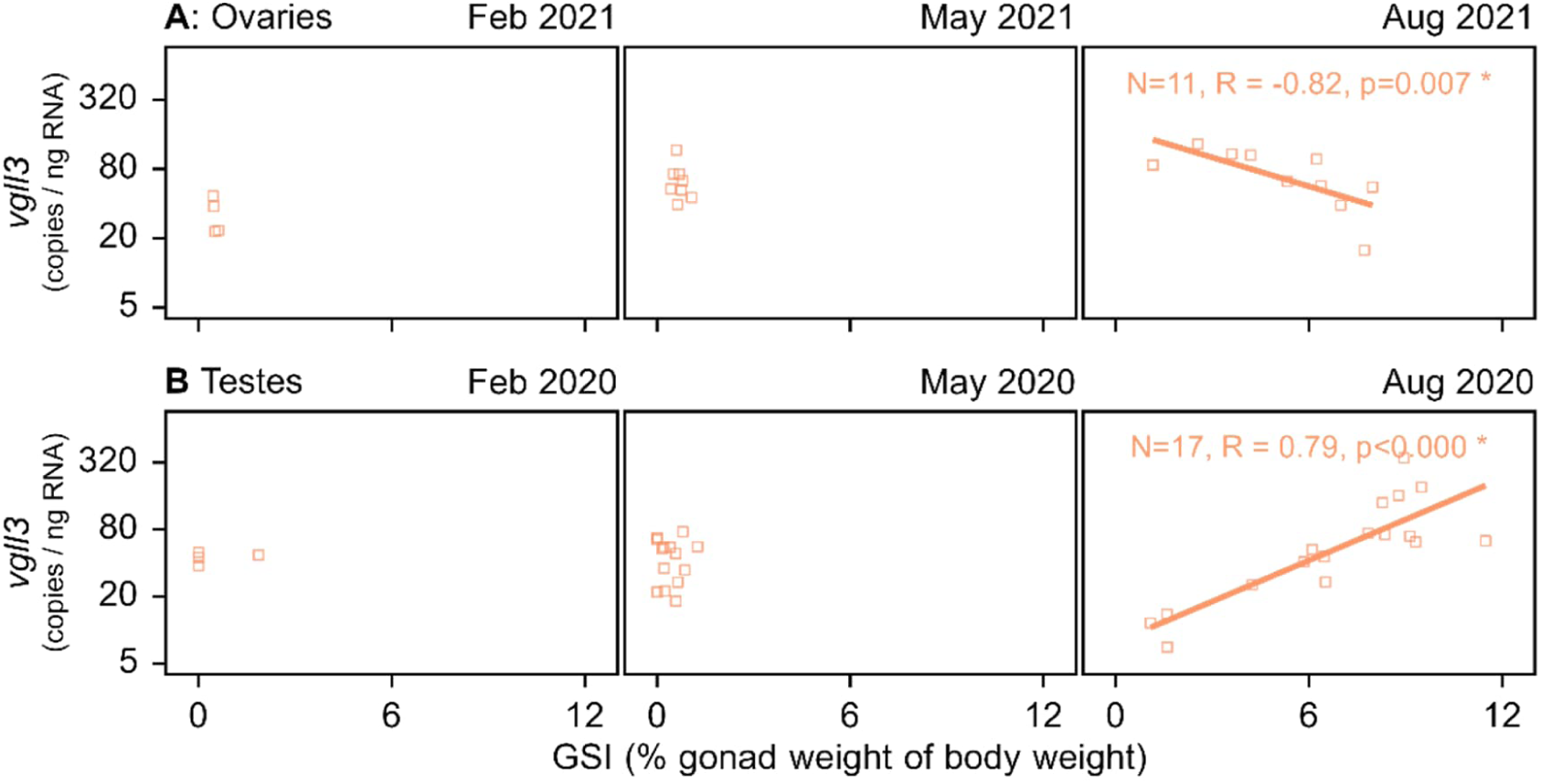
GSI and expression of *vgll3*. Relationship between gonadosomatic index (GSI, % gonad mass of whole body-mass) and gonadal *vgll3* expression (copies ng_-1_ RNA) in ovaries (A) and testes (B) at the three sampling time-points leading up to the second spawning season. Data are only shown for developing gonads (i.e., not fully mature, but showing some physical sign of development). Lines, shown only at the last time points, are for illustrative purposes and represent simple linear regression models of relationships between *vgll3* and GSI. N indicates sample size, R represents the degree of correlation (Spearman) and p the statistical significance. Asterisks (*) indicate p values below 0.05. The y-axes are log_e_ scaled.

The low-fat feed treatment, which started in July 2019, had a negative effect on body condition from August 2020 (Supplementary figure 3). Feed effects on *vgll3* expression were detected, but were dependent on time point and sex. In the first spawning season, there were no detectable effects of feed treatment on *vgll3* expression. However, in the period leading up to the second spawning season (as mentioned above), there was a significant effect of feed treatment on testicular *vgll3* expression. While there was an increase in *vgll3* expression over these three time points for the control-feed individuals, there was no statistically significant change over time for the low-fat treatment individuals. Specifically, for the low-fat treatment individuals, the estimated change in *vgll3* expression per month was 0.95-fold [0.80, 1.14]. Over the same time period (leading up to second spawning season), we detected 1.58 times higher [0.85, 2.80] expression of *vgll3* in the testes of control-feed treatment individuals, compared to the low-fat feed treatment. Given the 95% CI of this effect includes unity, there is some uncertainty about this effect, but 93.3% of the probability mass of this effect is above 1. Considering the following year (post second spawning season, leading up to the third spawning season), there were signs of a similar expression difference in testes between the feed treatments (Supplementary figure 3). In ovaries, no effects of the feed treatments on *vgll3* expression were detected.

### Expression of *amh* and *lhr*

Anti-müllerian hormone (encoded by the gene *amh*) is known to act as an inhibitor of spermatogenesis (Morais et al., 2017; Pfennig et al., 2015). Expression of *amh* is therefore a useful marker for maturation in testes, and we included *amh* expression measurements to gain additional context on the maturation process. As expected, the expression of *amh* was reduced in developing testes in late summer (August) in both years (Figure 4 B), but higher in winters. Ovarian *amh* expression showed clear signs of decreased expression only in fully mature ovaries (Figure 4 A). Interestingly, we detected positive correlations between expression of *vgll3* and *amh* in both testes and ovaries. However, these correlations depended strongly on season and developmental stage. In ovaries (Figure 5 A), these correlations were significant for undeveloped ovaries only in summer and autumn, for developing ovaries only in autumn, and never for mature ones. In testes (Figure 5 B), the correlations between *vgll3* and *amh* were significant for undeveloped testis in summer, autumn, and winter; for developing testes only in summer and autumn; and for mature testes in all seasons where they were present.

**Figure 4.**
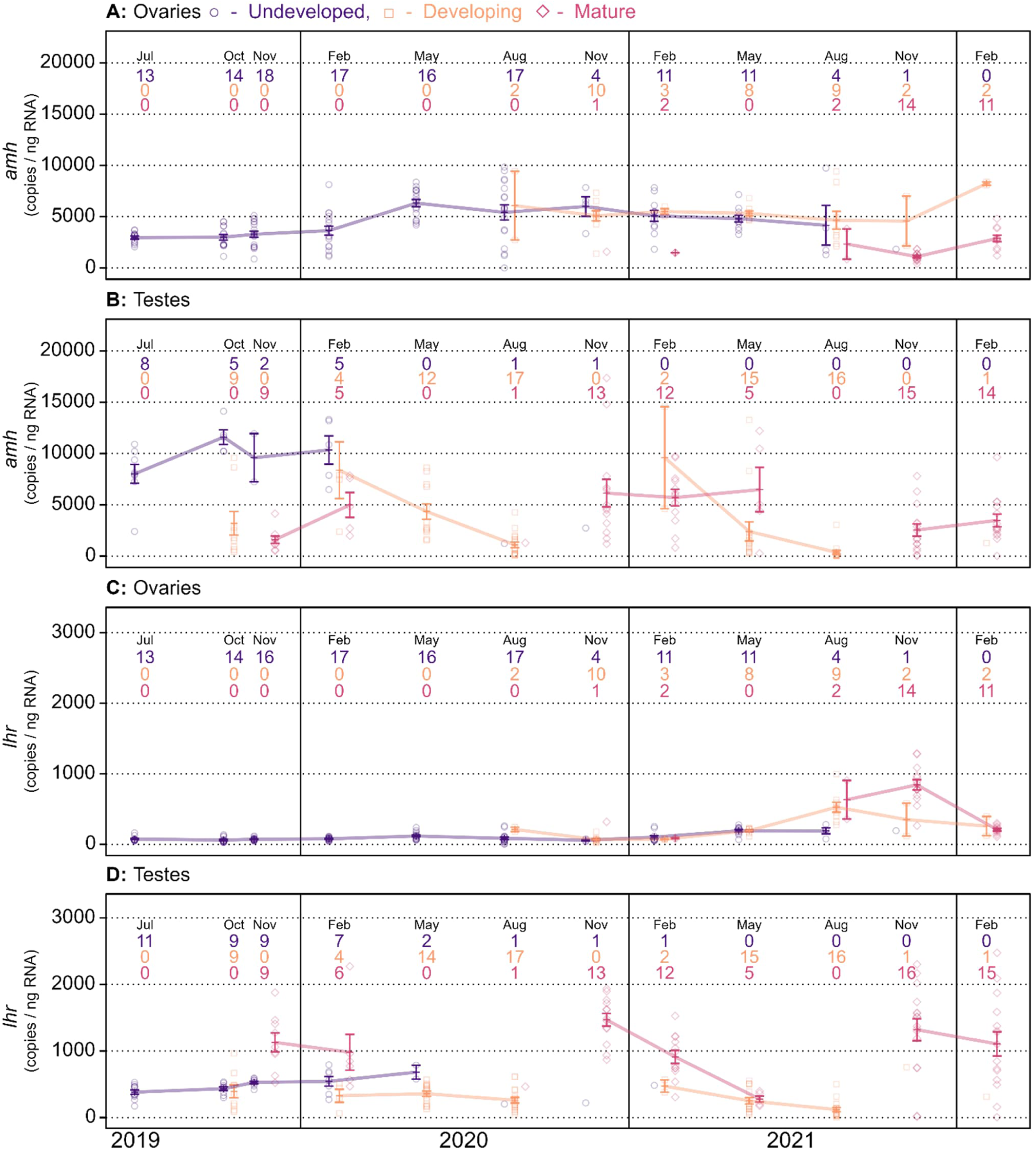
Gonadal maturation and *amh* and *lhr* expression over time. Gonadal expression of *amh* (A, B) and *lhr* (C, D) mRNA over the entire study period in ovaries (A, C) and testes (B, D), measured as copies ng_-1_ of RNA. Colours and shapes correspond to the maturation status of the gonads: purple circles for undeveloped gonads (no sign of enlargement), orange squares for developing gonads (some sign of enlargement), and red diamonds for fully mature gonads (releasing sperm or eggs). Lines connect means between time points with at least two samples and error bars indicate SE. The numbers on top indicate sample sizes for undeveloped, developing, and mature gonads. The y-axes are log_e_ scaled.

**Figure 5.**
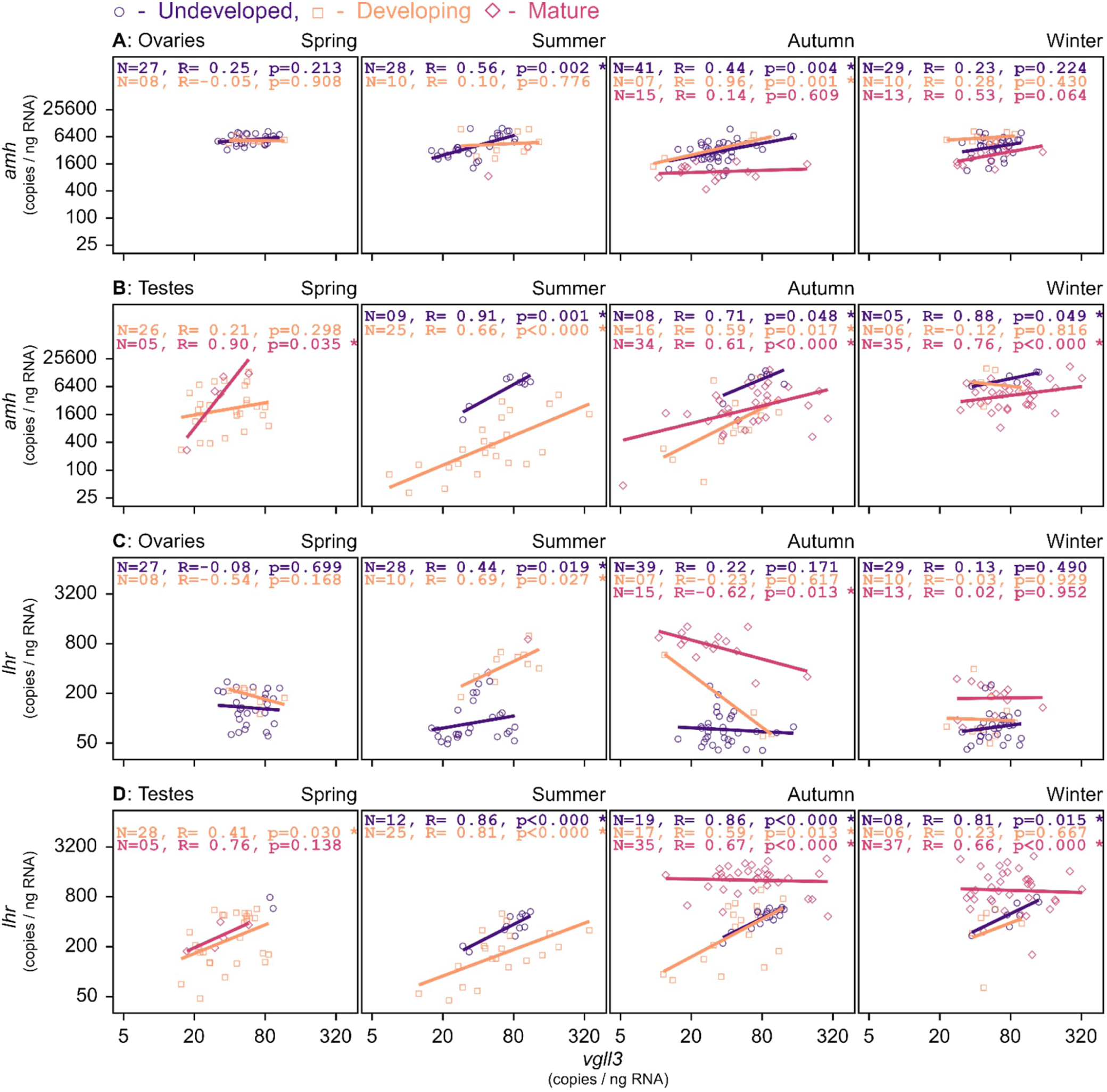
Correlated expression of vgll3, amh and lhr. Associations between gonadal RNA expression (copies ng_-1_ RNA) of *vgll3* and either *amh* (A, B) or *lhr* (C, D) in for different seasons (over all years) in ovaries (A, C) and testes (C, D) of different maturation status. Colours and shapes correspond to the maturation status of the gonads: purple circles for undeveloped gonads (no sign of enlargement), orange squares for developing gonads (some sign of enlargement), and red diamonds for fully mature gonads (releasing sperm or eggs). Lines, for illustrative purposes, represent simple linear regression models of relationships between *vgll3* expression and expression of either *amh* or *lhr*. N indicates sample size, R represents the degree of correlation (Pearson) and p the statistical significance for the groups of the same colour. Asterisks (*) indicate p < 0.05. The x- and y-axes are log_e_ scaled.

The luteinizing hormone receptor (encoded by the gene *lhr*) can be used as a marker of maturation in ovaries as its expression gradually increases towards maturation (Andersson et al., 2013). As expected, *lhr* expression increased towards maturation in ovaries and was highly expressed also in mature testes (Figure 4 C and D). Some significant correlations between expression of *vgll3* and *lhr* were detected. In ovaries (Figure 5 C), correlations between the expression of *vgll3* and *lhr* were more sporadic, and only present in summer for undeveloped and developing ovaries, and in autumn for mature ovaries. In testes (Figure 5 D), expression of *vgll3* and *lhr* correlated positively for undeveloped testes in summer, autumn, and winter; for developing testes in summer and autumn, and for mature testes in autumn and winter.

### Maturation, *vgll3* genotype and feed

The association between *vgll3* genotype and age at maturity is already well established, also for the males of the source experimental population used in this experiment (Åsheim et al., 2023). Despite the low sample sizes, we tested for a similar association in the individuals used in the current study. An association between maturation probability and *vgll3* genotype was indeed detected for males in the first spawning season, but not for females. In the second spawning season, almost all individuals matured regardless of genotype. In the first spawning season, for an average spring body weight, the mean predicted probability of having either developing or mature testes (vs. undeveloped) was 24.9% and 80.0% for a *vgll3*\*LL and *vgll3*\*EE individuals, respectively [95% CI of difference: 14.4, 79.7]. For females, there was no detectable effect of *vgll3* genotype on maturation probability in the first spawning season. Using the same comparison as above for females gave predicted probabilities of 49.1 and 48.9 % for *vgll3*\*LL and *vgll3*\*EE individuals [95% CI of difference: −47.3, 47.4]. We detected no significant effects of the feed treatments on maturation probability in the first spawning season for neither males nor females.

### Genotype and expression of *vgll3*

Studying expression variation between different *vgll3* genotypes within the maturation categories is challenging as there are already *vgll3*-maturation and maturation-expression associations present. To examine possible differences, we plotted *vgll3* expression over time for different *vgll3* genotypes separately for each maturation category (undeveloped and developing/mature) and sex (Supplementary figure 2). However, in both testes and ovaries, we observed that associations between *vgll3* genotype and expression were overall weak and differed between time points; no genotype (EE/EL/LL) was consistently expressed higher or lower than the others. Thus, over the full time series, we could not identify consistent patterns of associations between *vgll3* genotype and mRNA transcript abundance.

## DISCUSSION

We have presented for the first time a continuous time series of gonadal development and *vgll3* expression in both male and female Atlantic salmon over a 3-year time period, covering multiple spawning seasons. From these data, we identified several novel aspects of *vgll3* expression patterns during development and maturation.

### Higher *vgll3* expression in the ovaries and testes of individuals with delayed maturation

The observation that *vgll3* expression was higher in immature, undeveloped ovaries in the first spawning season suggests that the same pattern of *vgll3*-associated inhibition of maturation is present in both males and females. However, this pattern contrasts to what was found by Kjærner-Semb et al. (2018), who observed an increase in ovarian *vgll3* expression going from the oildrop prepubertal stage to the early pubertal stage. Compared to the expression pattern we found, the pattern observed in Kjærner-Semb et al. (2018) may correspond to the slight increase in *vgll3* expression we observed going from February to August after the first spawning season. In males, the association between testicular *vgll3* expression and maturation status has been suggested to be mediated via a *vgll3*-effect on the Hippo pathway. In this scenario, *vgll3* competes with Yap1/Taz for access to TEAD elements, affecting processes related to cellular differentiation (Kjærner-Semb et al., 2018; Kurko et al., 2020; Verta et al., 2020). The Hippo pathway has been found to associate with both ovarian and testicular development in other vertebrates (Clark et al., 2022; Kawamura et al., 2013; Lyu et al., 2016; Sen Sharma et al., 2019; Xiang et al., 2015), and has also been found to be involved in a range of other developmental processes (Meng et al., 2016). In any case, these results suggest that *vgll3* expression is similarly involved in the early maturation of males and females, at least in the sense that it involves a higher level of *vgll3* expression for immature individuals.

### Expression during development reflects inhibition of early testis development

Transitioning from the first to the second spawning season, we observed a change in the pattern of *vgll3* expression. For testes which were immature in the first spawning season, *vgll3* expression was high but then dropped in the following winter, slowly increasing up towards the next spawning season, while correlating positively with GSI as testis were gaining mass. Testicular development happens in sequential stages with regards to the proliferation and differentiation of spermatogonia and the Sertoli cells encysting them (Clermont, 1972; França et al., 2015; Schulz et al., 2010). Following this, we interpret from the expression patterns mentioned above that *vgll3* might be mainly involved in the inhibition of the earlier stages of testicular development. This mechanism was similarly hypothesized by Kjærner-Semb et al. (2018) on the grounds of histological observations as well as an observed downregulation of *vgll3* in maturing testes and a further increase in *vgll3* expression in regressing testis (post-spawning). Here, our data provide further support to this hypothesis by showing that *vgll3* expression is positively correlated with testis growth in later years, thus increasing not only after maturation (as in Kjærner-Semb et al., 2018) but also in the process leading up to maturation. Notably, for mature testes, the mean *vgll3* expression was low during the first spawning season, but high during the second one. This could indicate that the age at spawning affects how changes in expression of *vgll3* are timed, or that the influence of *vgll3* during spawning changes with age or testicular development. Taken together, we hypothesize from these results that that the earlier stages of testis development involve a downregulation of *vgll3* expression, while later stages involve a gradual upregulation of *vgll3*, and that the extent or timing of this upregulation may vary with age.

### Sex-specific expression dynamics

While we observed an increase in *vgll3* expression with GSI in testes, the pattern was more complex for ovaries. After the initial drop in *vgll3* expression as the ovaries started to develop, there was a steady increase in its expression, but then a negative association between GSI and *vgll3* expression in the autumn as the ovaries were starting to gain mass. These opposing patterns of *vgll3* expression and GSI between testes and ovaries could indicate that *vgll3* is involved in processes that are differently regulated in males and females. An example of such a process is estrogen (17β-oestradiol) signalling, which in males is high in the early stages of testis development, but in females coincides with the later stages of ovary development (Billard & Breton, 1978; Clelland & Peng, 2009; T. Miura et al., 2003). In testis, estrogen promotes spermatogonial germ cell- and Sertoli cell proliferation in the early phases of testis development (Amer et al., 2001; T. Miura et al., 1999, 2003; Song & Gutzeit, 2003). In ovaries, current knowledge of the role of estrogen is that it plays an important part during the vitellogenic phase of oogenesis (i.e., the phase of secondary oocyte growth)(Clelland & Peng, 2009). Estrogen may also be important at the earliest stages of oogenesis (C. Miura et al., 2007; See also Wootton & Smith, 2014, Ch 5).

Changes in estrogen signalling could involve both changes in estrogen production and changes in the expression of the estrogen receptor (esr1). In humans, expression of *VGLL3* has been negatively linked to expression of *ESR1*, the gene encoding the estrogen receptor ERα (Ma et al., 2022). Notably, ERα has been found to be central for the role of Sertoli cells during the first stage of testis development (spermatogonial stem cell renewal) (T. Miura et al., 2003). Production of estrogen itself, on the other hand, can be inhibited by *amh* via an inhibiting of cyp19a1 (aromatase, or estrogen synthase), an enzyme that converts testosterone to estrogen (Rodríguez-Marí et al., 2005). In this study, we found *vgll3* to have correlated expression with *amh*, so if one assumes a negative association between *vgll3* and estrogen signalling, then it can also be hypothesised that *amh* is involved in the process by inhibiting the testosterone-estrogen conversion. However, this hypothesis does not fit well with the observation that testicular *amh* expression was at its lowest at the same time as estrogen signalling should have been low (Billard & Breton, 1978). Changes in the levels of estrogen and testosterone could potentially also alter behaviours that are modulated by these hormones (See: Borg, 1994; Dey et al., 2010; Fernald, 1976). In this context, it is worth noting that behavioural trials have found a higher frequency of aggressive behaviour in juvenile individuals homozygous for the late *vgll3* genotype (Bangura et al., 2022).

While the hypothetical link between estrogen signalling and *vgll3* expression is intriguing, we underscore that it is at this time highly speculative, and that we include it as an example of how the results of this study could generate further hypotheses regarding the pathways *vgll3* is involved in.

### Expression and genotype of *vgll3*

To assess whether some of the effect of *vgll3* may be linked to genotype-specific expression levels, we examined whether any of the genotypes were more highly expressed than the others over the study period. This analysis was done after splitting the samples into groups of either undeveloped or developed/mature, accounting for a potential scenario where maturation status affects *vgll3* expression (reverse causality). Here, we found no significant and consistent association between genotype and expression. Previously, Verta et al. (2020), found a 14% higher expression of *vgll3* exon 2 (same exon as in this study) in *vgll3*\*LL individuals, but the variation was high with a highly overlapping range of expression between individuals of the two genotypes. Our sample sizes per genotype per time-point were overall low, so it is possible that we did not have sufficient statistical power to detect an effect, if it existed.

### Feed effects

The difference in *vgll3* expression between feed treatments, which was only observed in testes in the time periods leading up to the second and third spawning season, provides the first indication that nutritional intake can affect the temporal dynamics of *vgll3* expression. Two potential interpretations of this interaction are 1) the feed treatment could be affecting the speed of development of the testis, thus resulting in different *vgll3* expression levels when measured at the same time point, or 2) pathways regulating *vgll3* expression are integrating the nutritional or energetic state (e.g., lipid storage) of the fish. This mirrors previous findings where we observed an interaction between *vgll3* genotype and body condition in affecting maturation (Åsheim et al., 2023) as well as an analysis indicating that some of the effect of *vgll3* on maturation might be mediated via body condition (Debes et al., 2021). Interestingly, here, the effect of feed treatment on *vgll3* expression was only observed in males, indicating that the relationship between nutrient availability and *vgll3* expression might be sex specific.

### Conclusions

This study provides new insights into the temporal patterns of gonadal *vgll3* expression in Atlantic salmon throughout the maturation process, especially filling in knowledge-gaps related to maturation in females. We also demonstrate that differences in feed nutrient content can be reflected in the expression of *vgll3*, providing further insight on the connection between *vgll3*, resource acquisition and age at maturity. Our findings provide starting points for further research on the role of *vgll3* in the maturation process. For example, studies looking at the differences observed between males and females could target mechanisms with known opposite effects between sexes on maturation-related processes. Additionally, while we have uncovered novel patterns in whole-tissue expression of *vgll3*, further studies are necessary to identify underlying mechanisms for these patterns, such as changes in regulation of expression, or changes in proportions of various cell types during development. This could help in identifying new mechanisms related to the function of *vgll3*.

## Author contributions

Conceptualization: CRP, JPV, ERÅ, PVD

Data curation: ERÅ, AR, ID, MF, PVD

Formal analysis: ERÅ, PVD, JP

Funding acquisition: CRP

Investigation: ERÅ, AH, PL, AR, MF, ID, EPA, CRP

Methodology: JPV, PVD, CRP, ERÅ,

Project administration: CRP, JPV,

Resources: CRP, JE

Software: ERÅ, PVD,

Supervision: CRP, JPV,

Visualization: ERÅ,

Writing - original draft: ERÅ, JPV, CRP, PVD, EPA

Writing - review & editing: ERÅ, JPV, CRP, PVD, ID, MF, AR

## Funding

Funding for this study was provided by the Research Council of Finland (grant numbers 314254, 314255, 327255 and 342851), the University of Helsinki, and the European Research Council under the European Articles Union’s Horizon 2020 and Horizon Europe research and innovation programs (grant no. 742312 and 101054307). Views and opinions expressed are those of the author(s) only and do not necessarily reflect those of the European Union or the European Research Council Executive Agency. Neither the European Union nor the granting authority can be held responsible for them.

## Acknowledgements

We acknowledge Valeria Valanne, Katja Maamela, Markus Lauha, and Suvi Ikonen for helping with tissue dissections; Suvi Ikonen, Katja Maamela, Markus Lauha, Nikolai Piavchenko, Paul Bangura, Tiemen Jansen, Shadi Jansouz, Teemu Mäkinen, Jack O’Callagan, Anna Toikkanen, Ilke van Gesten, Pirta Palola, Fin Morrison, Jeferson Florez, Juho Kökkö, Matias Boehm, Mikko Immonen and Simon Lecarte for their assistance with animal husbandry and data collection; Jacqueline Moustakas-Verho, Kzenia Zueva, Marion Sinclair-Waters, Nico Lorenzen, Victoria Pritchard, Yann Czorlich, Suvi Ikonen, Spiros Papakostas and the staff at the Laukaa and Taivalkoski hatcheries for assistance with egg and sperm collection and/or fertilizations; Tutku Aykanat for calling SNPs; Seija Tillanen, and Shadi Jansouz for help with *vgll3* genotyping and sexing; Xindi Huang for discussions about ovary 5S rRNA variation; Susanna Airaksinen, Thomas Ginström Raisioaqua Ltd for producing the control- and low-fat feeds; John Loehr for assistance with logistics at Lammi Biological Station; and Natural Resources Institute Finland (LUKE) for access to their broodstocks.

## SUPPLEMENTARY MATERIALS

**Supplementary Table 1.**
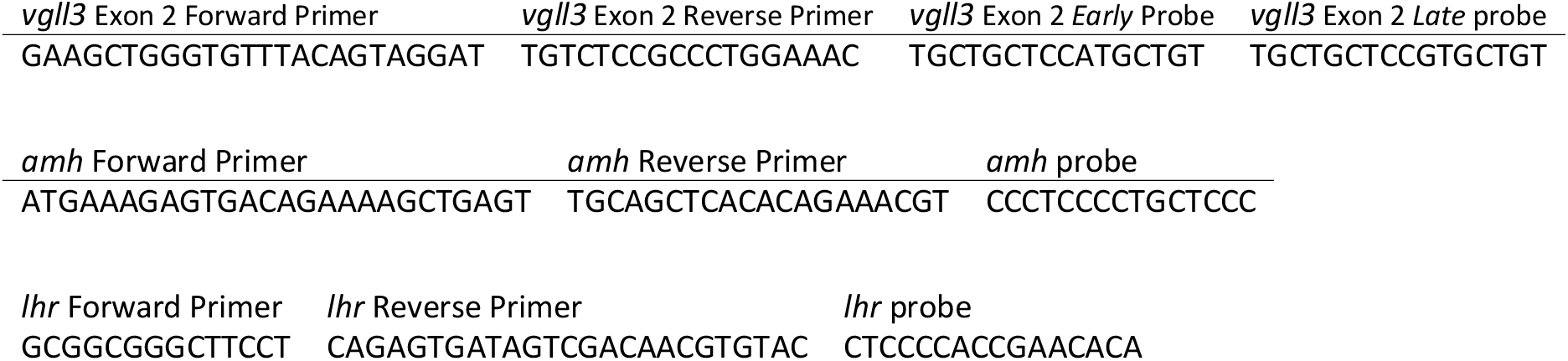
TaqMan assays.

**Supplementary figure 1.**
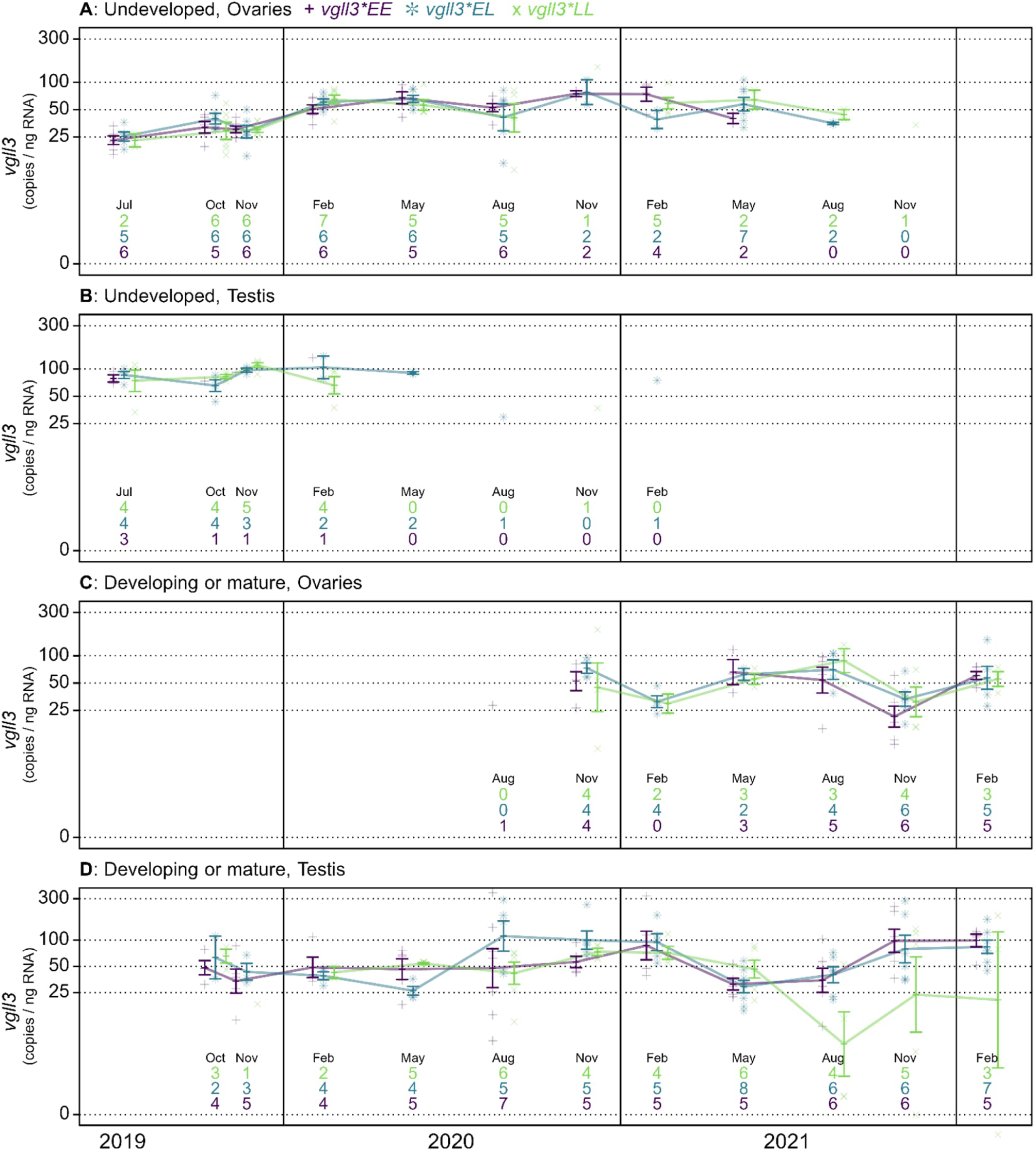
Gonadal *vgll3* genotype and expression over time. Expression of *vgll3* mRNA in gonads over the entire study period, measured as copies ng_-1_ of RNA, for all sampled gonads. Upper plots show values for undeveloped ovaries (A) and testes (B), and lower plots show values for developing or mature ovaries (A) and testes(B). Colours and shapes correspond to the genotype of the sampled individuals: purple (+) for *vgll3*\*EE, blue (❊) for *vgll3*\*EL, and green (x) for *vgll3*\*LL. Lines connect means between time-points with at least 2 samples and error bars indicate SE. The numbers on bottom indicate sample sizes for *vgll3* genotypes LL, EL, and EE. The y-axes are loge scaled.

**Supplementary figure 2.**
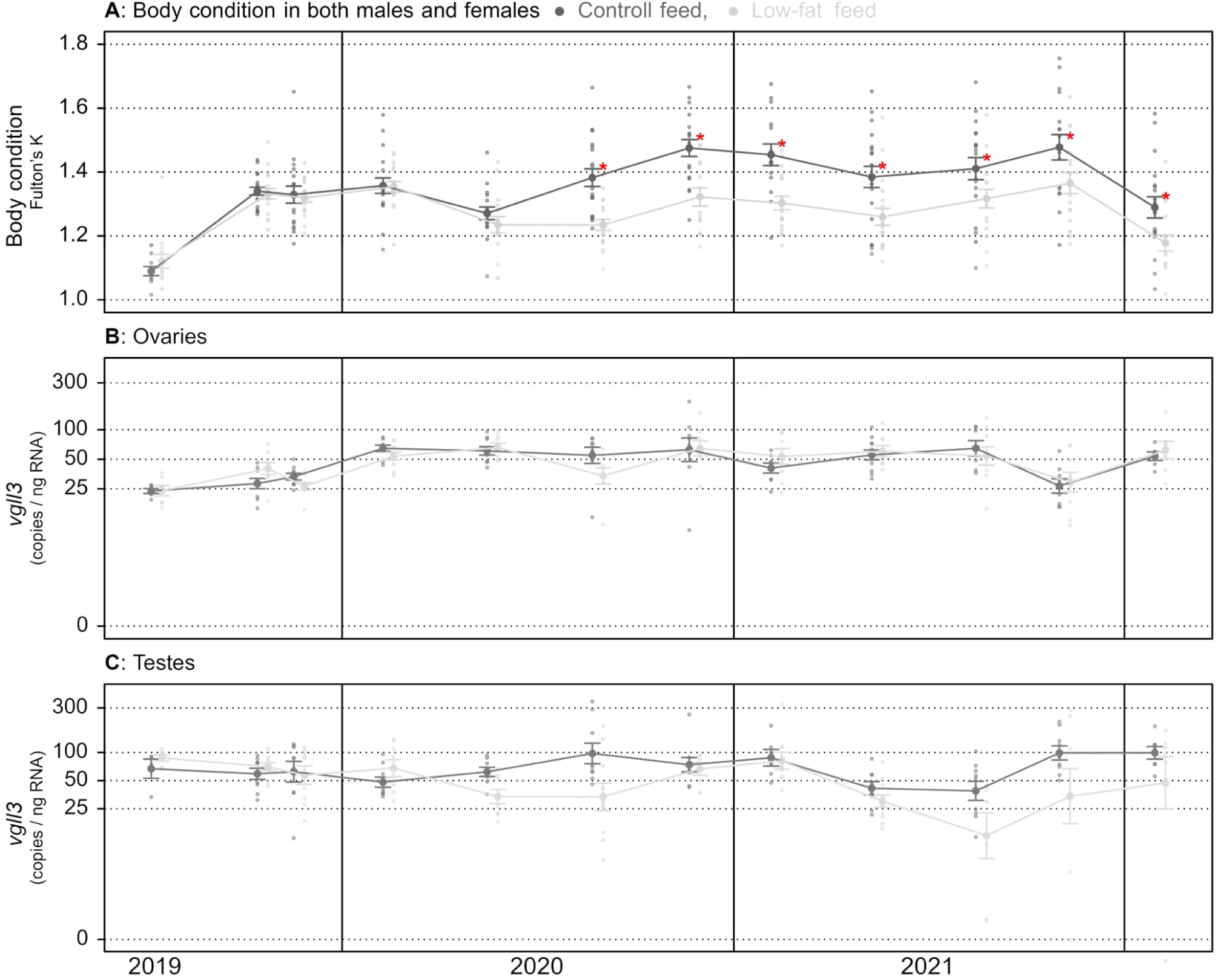
Body condition and *vgll3* expression over time, for two feed treatments. Body condition (A) and gonadal expression of *vgll3* mRNA (B: ovaries, C: testes) for the entire study period for all sampled individuals. Dark points represent individuals from the control feed treatment, and light grey points the low-fat feed treatment. Larger points and error bars indicate means and SE. Red stars in (A) indicate the significance (p > 0.05) of *t*-tests between the two treatments at the corresponding time point. The y-axes for B and C are log_e_ scaled.

**Supplementary figure 3.**
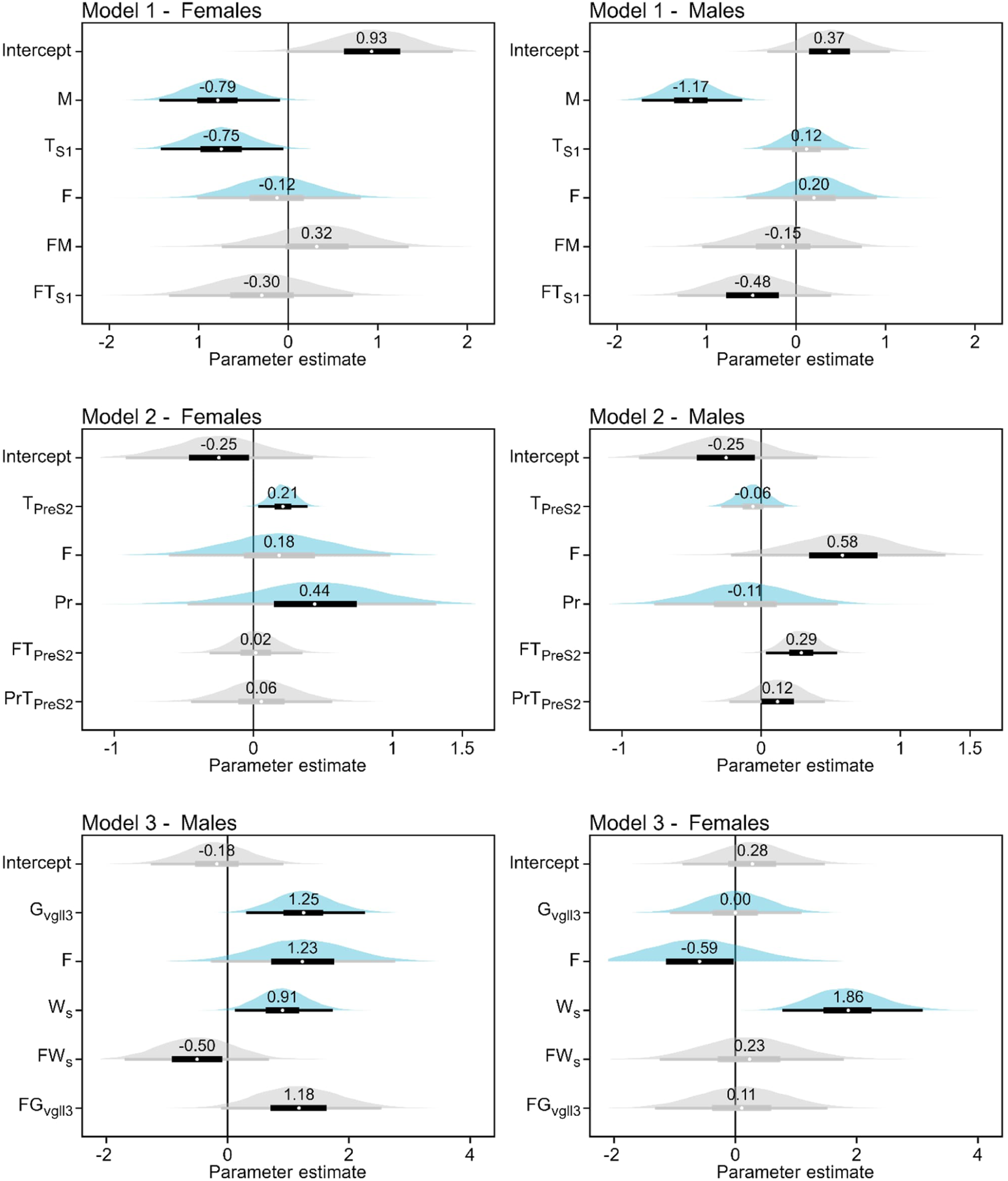
Posterior distributions of fixed-effect parameter estimates (*b_i_*) for all fixed-effects (*i*). See statistical methods for a descriptiption of model structures and parameters. Thick and thin bars indicate 50% and 97.5% credible intervals, respectively. Bars are colored gray if their intervals include zero. Blue distributions indicate that the parameter estimates are unconditional (marginal) of their non-significant interactions. Numbers indicate the mean parameter estiamte.

## Notes

### Competing Interest Statement

The authors have declared no competing interest.

### Summary of Updates

One co-author added to match the author list of the originally submitted pdf (accidental exclusion)

